# Bilateral auditory processing studied by selective cold-deactivation of cricket hearing organs

**DOI:** 10.1101/706085

**Authors:** Xinyang Zhang, Berthold Hedwig

**Affiliations:** Department of Zoology, University of Cambridge, Cambridge CB2 3EJ, United Kingdom

**Keywords:** Peltier element, chill coma, reciprocal inhibition, Omega neurons

## Abstract

We studied bilateral processing in the auditory ON neurons of crickets using reversible cold-deactivation of the hearing organs by means of Peltier elements. Intracellular recordings of the neurons’ activity in response to acoustic stimuli were obtained, while either the ipsilateral or the contralateral hearing organ was cold-deactivated. Afferent activity was abolished at a temperature of about 10°C. In ON1 contralateral inhibition has no effect on the latency and amplitude of the phasic onset activity, it enhances the decline of the onset activity and decreases the subsequent tonic spiking response to acoustic stimuli. As a consequence the phasic onset activity becomes more salient and reciprocal inhibition may support the detection of sound pulses. Contralateral inhibition had a significant impact on the tonic ON1 response, in line with its presumed function to enhance the bilateral auditory contrast. In ON2, experiments confirmed a bilateral excitatory input, with the ipsilateral input dominating the response, and no inhibitory coupling between the ON2 neurons.

## INTRODUCTION

For the analysis of bilateral auditory processing, insects like crickets and bush-crickets have been favourable systems (Lewis, 1983). Their hearing organs are positioned in the front legs, and are coupled by an auditory trachea (Michelsen et al., 1994, Schmidt and Römer, 2013) while auditory afferents terminate in the first thoracic ganglion (Esch et al., 1980), where they synapse to auditory interneurons (Popov et al., 1978, Wohlers and Huber, 1978, 1982, Hedwig and Stumpner, 2016). This arrangement allows for the selective decoupled stimulation of the hearing organs, while recording the activity of central auditory neurons, which provide the first stage of bilateral auditory processing, a crucial step for sound localization. In crickets bilateral auditory processing has been studied using decoupled stimulation of the hearing organs with miniature phones attached to the legs, while the central auditory trachea was severed (Kleindienst et al., 1981, Wiese and Eilts, 1985, Horseman and Huber, 1994a). Such an interference with the auditory system is not reversible, and as the auditory trachea is damaged it may also alter the function of the pressure difference receiver. Neural activity however, is highly temperature-dependent, a fact that in insects has been successfully used to study behaviour and neural processing (Farley et al. 1967, Matthews and White 2016). Cooling an insect leads to the loss of motor coordination at the critical thermal minimum and finally abolishes spike initiation and conduction in the nervous system when entering the chill coma (Overgaard and MacMillan, 2017; French, 1982, 1985). In bush-crickets and crickets the exposed position of the hearing organ allows to selectively cool and block the activity of auditory afferents without affecting the activity of the CNS (Baden and Hedwig, 2010). We further developed and refined this technique in order to analyze the dynamics of bilateral auditory processing in cricket ON neurons. We measured and compared their auditory responses under normal conditions and when one of the hearing organs was transiently deactivated by cooling. Two types of local omega-shaped neurons (ON1 and ON2) form bilateral mirror image pairs. Each ON1 receives excitatory inputs from the auditory afferents ipsilateral to its dendrites, and each ON1 neuron forwards inhibition to its contralateral sibling, so that both ON1 neurons are coupled by reciprocal inhibition (Selverston et al., 1985). The local circuit formed by the ON1 neurons is believed to enhance the bilateral auditory contrast and to support sound localization (Wohlers and Huber, 1978, 1982; Kleindienst et al., 1981). The ON1 network has also been suggested to contribute to auditory pattern recognition (Wiese and Eilts, 1985; Wiese and Eilts-Grimm, 1985) and may support the detection of species-specific sound pulses (Hedwig, 2016). The ON2 neuron has dendritic arborisations in both halves of the ganglion while its function is not yet clear.

## MATERIALS AND METHODS

### Experimental animals

Adult female crickets (*Gryllus bimaculatus* DeGeer) from a colony at the Department of Zoology/Cambridge were used 10-25 days after final ecdysis and were checked for physical integrity of their tympanic membranes. Experiments were performed at 24°C room temperature.

### Peltier elements for cold-deactivation of hearing organs

Two 10×10 mm Peltier elements (Peltron, Fürth, Germany) were used to control the temperature of the front tibiae and inactivate the ears. The Peltier devices were placed on both sides of the cricket at a tilted angle of 45°. The front tibiae were attached with Plasticine to the cooled surface of the elements, the heated side was attached to an 80×80×50 mm aluminium block acting as a heatsink (Fig. 1A,B). Crickets were mounted ventral side up on the centre of a platform covered by Plasticine, avoiding electrical conduction between the animal and the Peltier devices. Thermocouples (K-PTFE thermocouples, RS components, Corby, Northants, UK) were calibrated and attached to the surface of the Peltier elements to monitor the surface temperature by a custom build circuit.

**Fig. 1A:**
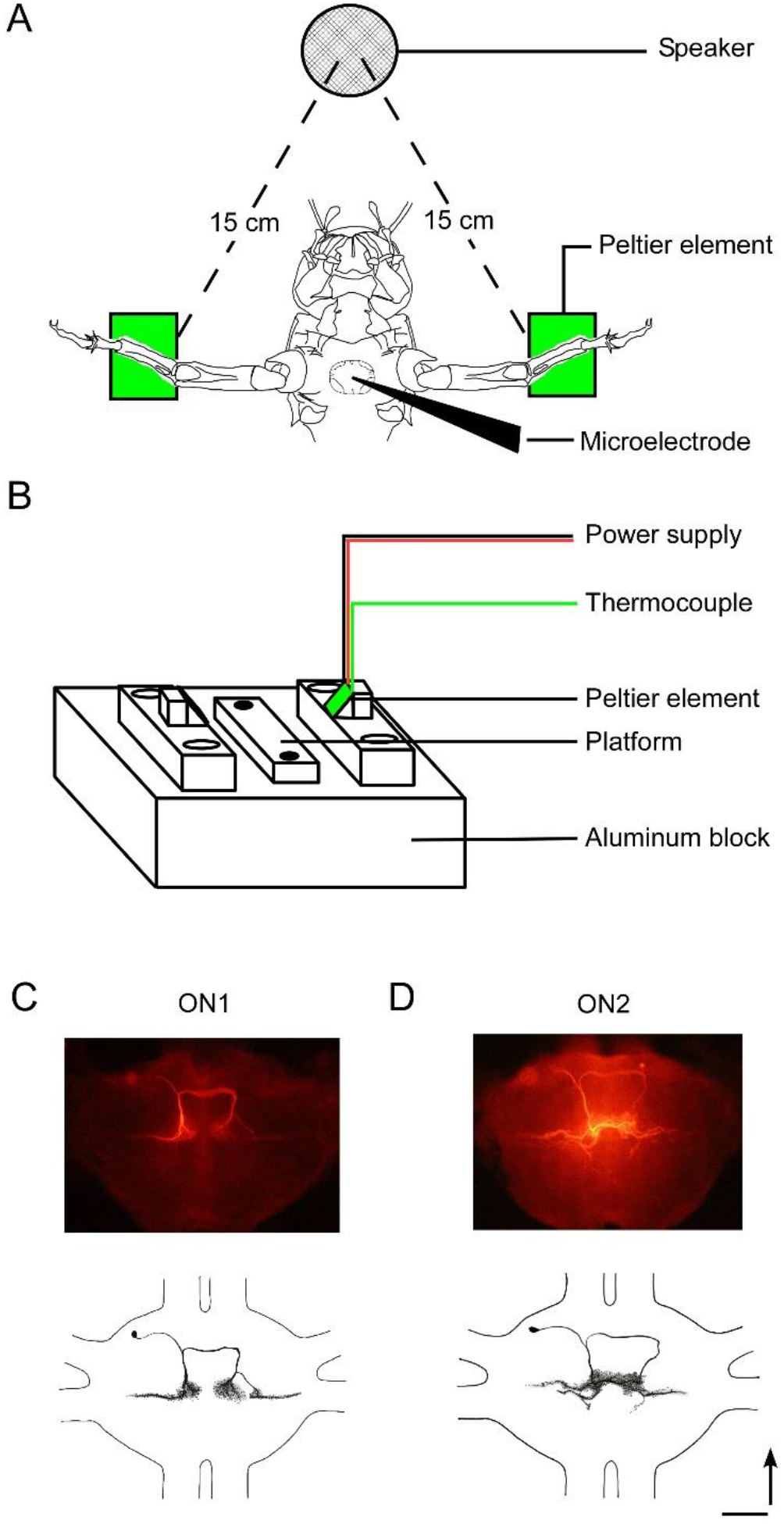
Experimental set up for cooling hearing organs and intracellular recordings of central neurons. (A) The cricket front legs are tethered to Peltier elements, the prothoracic ganglion is exposed for intracellular recordings, and a loudspeaker is placed in front of the animal for acoustic stimulation. (B) Details of experimental device with the platform to mount the cricket, the arrangement of the Peltier elements and the thermocouples, and the aluminum block serving as heat sink. (C, D) Structure of ON1 and ON2, reconstructed from image stacks after labelling with Alexa 568.

### Acoustic stimulation

Sound pulses of 150 ms duration, 250 ms inter-pulse intervals, a carrier frequency of 4.8 kHz, and an intensity of 75 dB SPL rel. 20 μPa were used. The rising and falling ramps for each pulse were 2 ms. Stimuli were designed with audio software (Cool Edit 2000, Syntrillium, Phoenix, AZ, USA) and were presented by a loud speaker (Sinus live NEO 13s, Conrad Electronics, Hirschau, Germany) placed in front of the specimen at a distance of 15 cm from each tympanic membrane (Fig. 1A). Intensity was calibrated at the location of the cricket using a measuring amplifier and a 1/2 inch free field microphone (Models 2610 and 4939, respectively; Brüel & Kjær, Nærum, Denmark).

### Intracellular recordings

The prothoracic ganglion was exposed (Fig. 1A), the trachea supplying the ganglion and the auditory trachea connecting the left and right ear were left intact, the mid and hind legs and the gut were removed. The ganglion was covered with insect saline; composition in mmol/l: NaCl 140; KCl 10; CaCl_2_ 7; NaHCO_3_ 8; MgCl_2_ 1; TES 5; D-trehalose dehydrate 4, adjusted to pH 7.4. A stainless steel platform with an optic fibre embedded was placed under the dorsal side of the ganglion for support and illumination, the platform also served as the reference electrode. To increase mechanical stability, a tungsten ring was gently placed on the ventral side of the ganglion. If crickets generated intense motor activity, the anterior connectives of the prothoracic ganglion were squeezed or cut.

Microelectrodes were pulled from borosilicate glass capillaries (Harvard Apparatus Ltd., UK; 1 mm OD, 0.58 mm ID) using a DMZ-Universal micropipette puller (Zeitz Instruments, Martinsried, Germany), and were filled with 2 M potassium acetate providing resistances of 40-60 MΩ. For iontophoretic staining, the tips were filled with 0.25% Alexa 568 hydrazide (Invitrogen, Thermo Fisher, UK) dissolved in water, the shaft was backfilled with 2 M KAc, giving a resistance of 80-130 MΩ. Neurons were recorded in their main dendrites. The position of the microelectrode was controlled by a Leitz micromanipulator (Model M; Leica Microsystems, Wetzlar, Germany). Electrode depth was monitored with a digimatic indicator (ID-C125MB; Mitutoyo Corporation, Japan) attached to the micromanipulator. The major arborisations of the Omega neurons were encountered at a depth of 150-200 μm. Intracellular recordings lasted from 10-60 min, signals were amplified by a DC amplifier (BA-01X, NPI Electronic, Tamm, Germany).

All recording channels (intracellular recordings, iontophoretic current, thermocouples, sound) were sampled at 20 kHz using a CED 1401 data acquisition interface (Micro1401 mk II, CED, Cambridge, UK), and saved with a PC for off-line analysis.

Experiments focussed on the ON1 and ON2 neurons in the prothoracic ganglion. Fluorescent dyes were iontophoretically injected into the neurons for 5-20 min by hyperpolarizing current of 1.5-3.0 nA. The tissue was processed with standard histological techniques. ON1 and ON2 morphology was revealed with an epifluorescence microscope (Axiophot, Carl Zeiss, Wetzlar, Germany) and was reconstructed from image stacks (Fig. 1C,D). Neurons were identified according to their morphology and response patterns (Watson and Hardt, 1996; Wohlers and Huber, 1982).

### Experimental design and data analysis

Neuronal activity was recorded in three different conditions: (1) Normal situation with both ears functional; (2) cold inactivation of the contralateral ear; (3) cold inactivation of the ipsilateral ear. We use *ipsilateral* and *contralateral* in respect to the cell body/main dendrite of the recorded ON.

For the auditory responses we calculated Peri-Stimulus-Time histograms; the latency between the onset of a sound pulse and the first spike of the response; the instantaneous spike rate based on the time interval between subsequent spikes, and the instantaneous spike-rate averaged over the set of stimuli. The auditory response showed a phasic onset peak; the duration of the peak was defined as the time from the start of the first spike to the point where the spike rate declined to 75% of the peak spike rate. The tonic response component was measured over a time window from 60-100 ms after the response onset. Recorded data were analysed with Neurolab software (Knepper and Hedwig, 1997) and Spike 2 (CED). Data are given as means and SEM.

## RESULTS

### Evaluating the effectiveness of the cooling system

We tested the effectiveness of the Peltier elements while recording an ON1 neuron intracellularly and simultaneously cooling the ipsilateral ear (Fig. 2). Activating the Peltier element decreased the temperature of the tibia from 24°C room temperature to 10°C within 60 s. When the Peltier element was turned off the tibia recovered to 24°C within 90 s. Before cooling ON1 responded to a sound pulse of 150 ms/75 dB SPL with a depolarization and about 25 spikes (Fig. 2A). With the decrease in temperature ON1 activity declined. At 90 s in the cooling process with the tibia at 10°C, sound evoked spike activity was abolished and the inhibition mediated by the contralateral ON1 neuron became obvious (Fig. 2B,C). Spikes still occurred in the interval between sound pulses, demonstrating that cooling the tibia did not affect ON1’s ability to generate spikes. When the tibia temperature recovered, ON1’s response reached the initial level again (Fig. 2 D,E); before cooling it was 24.9 ± 13.4 AP/pulse and 2 min after cooling 24.9 ± 12.4 AP/pulse (*P* = 0.99) (Fig. 2A,E). Allowing for some variation in animals and experiments we subsequently set the cooling temperature to 8-10°C to transiently block the activity of one or both hearing organs.

**Fig. 2:**
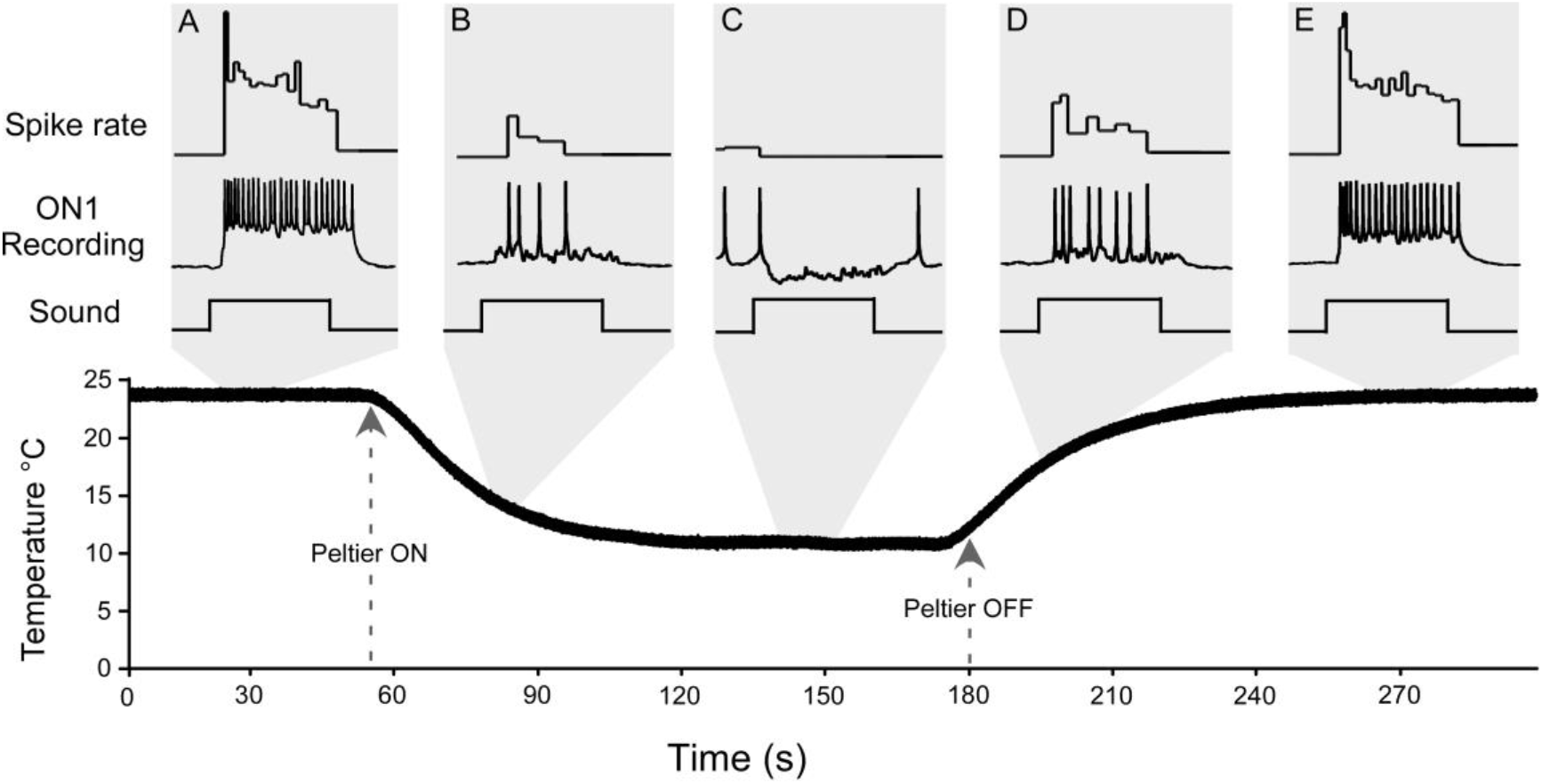
ON1 response to sound pulses during cooling and recovery of the ipsilateral hearing organ. (A) The normal excitatory input to ON1. (B,C) With the Peltier device switched *on* the ipsilateral hearing organ is cooled to 10 °C within 90 s, excitation to the ON1 neuron is abolished and the contralateral inhibition dominates its response. (D,E) With the Peltier device *off*, the ON1 excitatory response recovers as the hearing organ reaches room temperature.

### Activity of ON1 in normal condition and when one ear is cold-deactivated

Intracellular labelling confirmed the typical shape of ON1 extending over both halves of the prothoracic ganglion with dendrites at the side of the cell body and axonal arborisations at the opposite side (Fig. 1C). When both ears were functional, ON1 responded to 150 ms sound pulses with a salient phaso-tonic spike pattern (Fig. 3A). In the example given, the brief onset peak reached a maximum firing rate of 385.9 Hz and lasted for 10 ms; the firing rate then declined to a tonic level of 116.6 Hz, which was maintained for the rest of the response. ON1 activity however, was variable in different preparations, the averaged results for all recordings (N=6) show an activity pattern with a peak rate of 268.2±32.9Hz and a tonic rate of 111.6±10.8 Hz.

**Fig 3:**
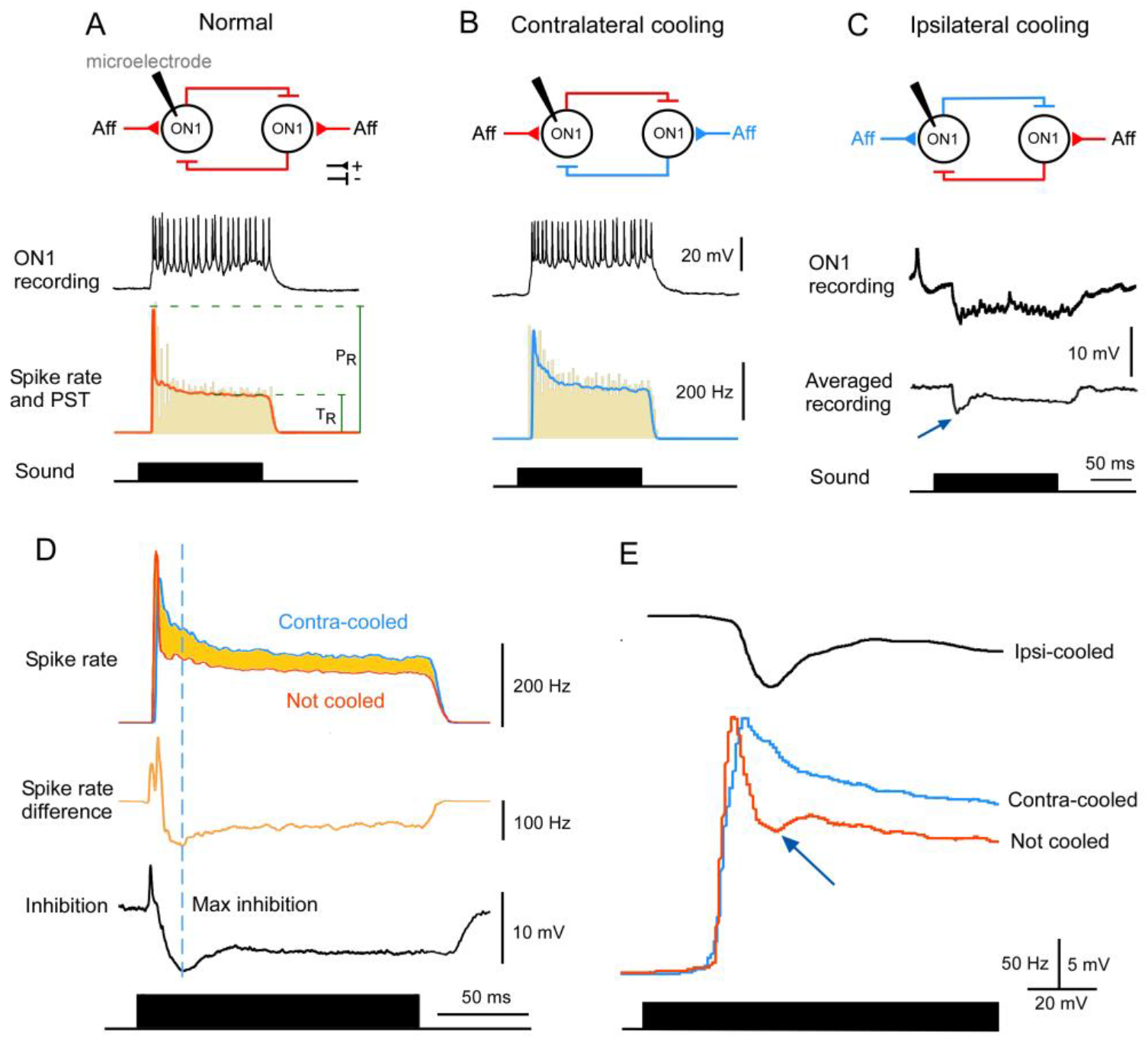
ON1 activity in three experimental conditions. (A) Normal, (B) contralateral ear cooled, and (C) ipsilateral ear cooled. (Top) Layout of active (red) and functionally blocked (blue) ON1 circuitry. (Middle) ON1 intracellular recorded response. (Bottom) Averaged ON1 instantaneous spike rate (red and blue) and the peri-stimulus time (PST) histogram (yellow) (n=192, 19±0.9 AP/pulse; n=179, 24.1±0.9 AP/pulse), P_R_: Peak spike rate, T_R_: Tonic spike rate. (D, Top): Averaged ON1 spike rate of recording in A and B under normal (red) and contralateral cooled (blue) conditions overlaid. (Middle) Difference in ON1 spike rate between the normal response and the response during contralateral cooling (N=14). (Bottom) ON1 response to contralateral inhibition when the ipsilateral ear is cooled (N=14). (E) Timing of the contralateral inhibition (top, black), the spike rate upon removing the contralateral inhibition (middle, blue) and the spike rate during the normal response (bottom, red). Contra lateral inhibition coincides with the fast decline of the onset peak spike rate and the subsequent “dimple” in the discharge rate.

When the contralateral ear was cooled, the reciprocal inhibition was functionally removed as the excitatory afferent input to the contralateral ON1 was abolished. In response to 150 ms sound pulses the same ON1 generated a phaso-tonic spike pattern, the phasic peak now lasted for 15 ms and reached a spike rate of 319.6 Hz, while the tonic activity level settled at 149.3 Hz. (Fig. 3B). The average peak and tonic activity of all recordings (N=6) were 252.6±22.5Hz and 142.12±8.9 Hz, respectively, and the ON1 tonic activity was now significantly higher than in the normal situation (111.6±10.8 vs. 142.1±8.9 *P*=0.0025). The peak spike rate however, was not changed by the contralateral inhibition (268.2±32.9 vs. 252.6±22.5, *P*=0.19).

The ON1 response substantially changed when we deactivated the ipsilateral auditory organ and abolished its excitatory afferent input. The ON1 activity now was determined by the inhibitory input from its contralateral sibling (Fig. 3C). In the example recording, an initial drop of the membrane potential reached a maximum of −9.5 mV relative to the resting membrane potential. The inhibition then decreased to −4.7 mV and remained at this level for the duration of the stimulus. On average (n=6) the initial maximum hyperpolarization mediated by the contralateral input reached −6.1±1.9 mV (Fig. 3C, arrowhead) and then declined to a constant level of −4.5±1.8 mV until the end of the stimulus.

### Impact of contralateral inhibition on ON1 activity

Removing functionally the inhibition from the contralateral ON1 allowed us to analyze the dynamic impact of the inhibition. The contralateral inhibition did not affect the latency of ON1, which was 18.51±0.53 ms in the normal situation and 19.50±0.59 ms when the contralateral ear was cooled (*P*=0.06, Fig. 3A,B). Also the timing of the phasic onset spike rate was not significantly different in both situations with normal: 21.09±0.60 ms and contralateral cooled: 22.92±0.64 ms (*P*=0.058). The average latency of the contralateral inhibition arriving at the recorded ON1 was 21.88 ±0.85 ms and was about 3.52±0.95 ms longer than the latency of the excitatory activity. The maximum of the inhibition was reached at 33.1±1.76 ms.

The impact of the ON1 reciprocal inhibition should be reflected in the spike rate of the auditory response. We overlaid the averaged instantaneous spike rate in response to a 150 ms sound pulse for the normal and the contralateral-cooled situations.

Subsequently we subtracted the normal spike rate from the spike rate obtained during contralateral cooling, demonstrating the effect of the reciprocal inhibition, as shown in Fig. 3D. For 14 recordings we calculated the spike rate difference which was maximum with 107.95±8.20 Hz after 32.4±1.70 ms and then reached a tonic level of 58.87±0.04 Hz. The time course of the spike rate difference showed a high similarity with the average time course of the membrane potential change caused by the contralateral inhibition when the ipsilateral ear was cooled (Fig. 3D bottom). The difference in the spike rate followed the course of the inhibition, both reached a peak at either 33.10±1.76 ms or 32.4±1.70 ms (r=0.65) and then settled at a tonic level.

### Does reciprocal inhibition contribute to pattern recognition?

For the same recordings we also analyzed if reciprocal inhibition could make the response to sound pulses of the cricket song more salient. Sound pulses of the calling song are about 17-20 ms long and come with a corresponding interpulse interval duration.

Analyzing the spike rate dynamics of ON1 neurons under normal conditions, revealed that following the phasic onset response 71% of the neurons exhibited an additional transient decrease in spike rate, which was beyond the expected decrease based on the time course of the decay. This transient decrease formed a pronounced ‘dimple’ in the decaying spike rate (Fig. 3E, arrowhead). When the inhibitory input was abolished, in experiments when the contralateral ear was cooled, the spike rate of the onset peak decreased more gradually and the dimple was not observed (Fig. 3E, contra-cooled). Recordings when the ipsilateral hearing organ was deactivated show that the contralateral inhibition reached its maximum at 33.10±1.76 ms (Fig. 3E, ipsi-cooled) while the ‘dimple’ in the spike rate reached its lowest point at 32.15±0.02 ms (r=0.96), indicating that the contralateral inhibition contributes to sharpening the initial phasic onset response of the ON1 neuron. On average the dimple ends at 38.92 ±2.71 ms, thus it would be well timed to reduce or suppress any spike activity in the interpulse intervals of the male calling song. Also the duration of the phasic onset response in normal conditions was significantly shorter than during contralateral cooling (10.11±1.11 ms vs. 15.32±1.54 ms, *P*=0.003).

We analyzed to what degree contralateral inhibition would have an effect on the relative representation of the sound pulses in the spike rates and analyzed the ratio of the phasic onset spike rate (P_R_) to the tonic rate (T_R_) (Fig. 3, see P_R_ and T_R_). This was 2.40±0.18 in the normal situation and as the tonic activity increased when the contralateral inhibition was removed the ratio dropped to 1.77±0.08. Thus, for the stimuli tested in normal conditions contralateral inhibition makes the onset activity significantly more pronounced relative to the tonic part of the response (*P*<0.005).

Taking all ON1 recordings the amplitude of the maximum inhibition and the ratio of the phasic/tonic spike rate between normal and contralateral cooled situations was well correlated (r=0.67), implying that a stronger inhibition increases the ratio between the initial peak spike rate and the tonic rate. In the context of song pattern processing this indicates that reciprocal inhibition sharpens the onset response of ON1 neurons to species-specific sound pulses, and that immediately after the processing of a normal sound pulse it may contribute to reduce the probability of spikes occurring in the interpulse intervals.

### Activity of ON2 in normal condition, and when the ipsilateral or contralateral hearing organ is cold-deactivated

ON2 neurons were recorded in the same area as ON1 and their typical structure with broad arborisations connecting both sides of the ganglion was revealed (Fig. 1D).

In the normal situation, the initial peak spike rate of the neuron (Fig. 4) was 337.7 Hz which declined within 15.5 ms to a tonic level of 167.9 Hz. When averaged across 5 ON2 neurons, the latency for the first spike was 19.1±0.5 ms, and the peak spike rate reached 308.9±50.6 Hz. Subsequently, the spike rate settled at a tonic level of 176.6±27.7 Hz. The pooled P_R_/T_R_ ratio of all ON2 neurons recorded in normal and contralateral cooled conditions was 1.76±0.14 and 1.75±0.16 (*P*=0.46), respectively.

**Fig 4:**
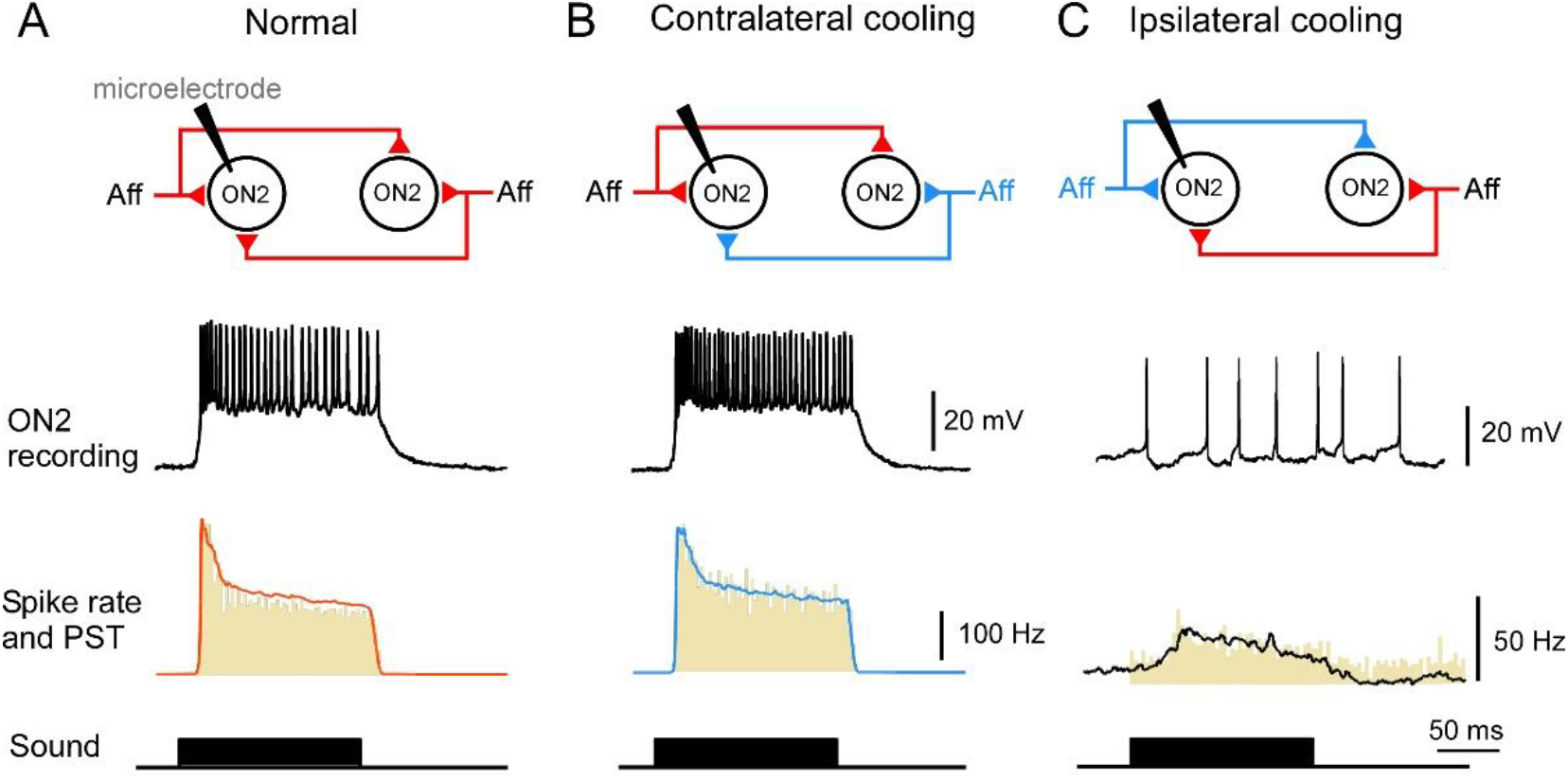
ON2 activity in three experimental conditions. (A) Normal, (B) contralateral ear cooled, and (C) ipsilateral ear cooled. (Top) Layout of active (red) and functionally blocked by cooling (blue) ON2 circuitry. (Middle) ON2 intracellular recorded response. (Bottom) Averaged ON2 instantaneous spike rate overlaid on PST-histogram (yellow), (A: n=432, AP/stim 28.5± 1.0; B: n=91, 35.2± 0.7, C: n=159, 3.7±2.4).

As the ON2 neurons are not indicated to be coupled by reciprocal inhibition (Wohlers and Huber, 1982) different effects were expected in the cooling experiments. In the contralateral-cooled condition, the example ON2 showed a similar response as in the normal situation (Fig. 4B). It generated a phasic onset spike rate of 322.0 Hz, which then declined to a tonic level of 179.5 Hz. When averaged over 5 ON2 recordings, the spike latency was 20.4±1.2 ms. The peak onset activity was 284.4±70.2 Hz, it occurred after 23.4±1.5 ms and then decreased to a tonic level of 169.2±42.3 Hz. Average spike latency in the normal and contralateral cooled experiments were 19.1±0.5ms and 20.4±1.2ms, respectively, and were not significantly different (*P*=0.24), also the peak spike rates in the normal and the contralateral-cooled situation were not significantly different (*P*=0.44). The inactivation of the ipsilateral ear led to reduced depolarization and a reduction in spike rate by 87.56 ±1.52% from in response to the acoustic stimulus as it received an excitatory synaptic input from the contralateral ear (Fig. 4C). The pooled P_R_/T_R_ ratio for normal and contralateral cooled experiments are 1.76±0.14 and 1.75±0.16 respectively (*P*=0.46). The experiments demonstrate a clear asymmetry between the bilateral excitatory auditory inputs, with the ipsilateral input dominating ON2’s response. An inhibitory input was not never observed.

## DISCUSSION

The excitatory ON1 auditory response failed when the ipsilateral fore-tibia was cooled to 10°C, demonstrating that low temperature abolishes activity of the auditory afferents. Our results are in line with French (1982, 1985) giving a critical temperature of less than 10°C for the deactivation of cockroach mechanosensory neurons and Baden and Hedwig (2010) reporting 4°C surface temperature of the Peltier device to block the afferents in a considerably larger bush-cricket. We speculate that for the cricket auditory afferents the cold-block temperature was between the critical thermal minimum temperature and their chill coma temperature, which in insects abolishes transduction and spike activity. The functional consequences of the cold-block may be linked to decreased membrane fluidity and/or altered ion channel kinetics which are reversible (Overgaard and MacMillan, 2016). Temperature could also affect the vibrations of the tympanic membrane, however in locusts a temperature shift from 21°C to 28°C had only a minor effect (Eberhard et al., 2014). Reducing the temperature of an insect appendage by Peltier elements is an effective approach to reversibly block inputs from peripheral sensory organs in order to analyze central neural processing. The method is suitable to block sensory afferents of insect appendages without surgical interference and should be versatile to study e.g. mechanoreceptive feedback involved in walking or jumping in larger insects (Burrows, 1996).

### Does reciprocal inhibition in ON1 contribute to auditory contrast enhancement and temporal filtering?

As the cold-deactivation of the auditory afferents was reversible, we could analyze in continuous intracellular recordings the function of bilateral auditory processing in ON neurons. For ON2, our experiments confirmed bilateral excitatory auditory inputs, with the ipsilateral input very much dominating the neuron’s response akin to recordings by Wohlers and Huber (1982). For ON1 neurons, we demonstrated that the time course of the contralateral inhibition matched the difference in spike rate with and without the inhibitory input. Contralateral inhibition had a significant impact on the tonic ON1 response, and this is in line with the presumed function of ON1 neurons to enhance the bilateral auditory contrast (Wohlers and Huber, 1978, 1982, Kleindienst et al., 1981, Wiese and Eilts, 1985). Reciprocal inhibition however, did not alter the response latency as also reported by Kleindienst et al. (1981) and the phasic onset spike activity. This is surprising as it indicates that contrast enhancement – at least at the level of the ON1 neurons – would not be effective at the level of bilateral spike timing. A high phasic onset activity related to the pulse pattern, may be relevant for controlling the activity of the ascending auditory neurons AN1 (Horseman and Huber, 1994a,b).

The ON1 results demonstrate that reciprocal inhibition can contribute to the temporal filtering of the calling song pattern by sharpening the phasic onset response (Hedwig, 2016). With contralateral inhibition a more pronounced decrease of the phasic response occurred as compared to a more gradual decrease when contralateral inhibition was blocked (Fig. 3C). Also a Ca^2+^ triggered potassium outward current (Baden and Hedwig, 2009; Sobel and Tank, 1994) could contribute to the rapid decrease of the onset spike activity. Contralateral inhibition however, plays the dominant role as the pronounced decline in spike rate was not observed when the contralateral ear was cold-deactivated.

The higher P_R_/T_R_ ratio of the auditory response with contralateral inhibition also indicates that the inhibition enhanced the coding of the onset of a sound pulse relative to the tonic response level. A salient onset response of ON1 will increase the signal-to-noise ratio and may reflect an adaptation of the cricket’s afferent auditory pathway to efficiently copy the species-specific sound pattern (Nabatiyan et al., 2003, Hedwig, 2016). The effect of contralateral inhibition in shaping the ON1 response will be strongest in a frontal stimulus situation, when both hearing organs are activated to the same extend, supporting Marsat and Pollack’s (2005) suggestion of a direction sensitive pulse filter mechanism.

Under the impact of contralateral inhibition in 71% of animals a decrease of the spike rate occurred forming a relative minimum, a “dimple” immediately after the phasic onset activity. This dimple coincided with the maximum of the contralateral inhibition (Fig. 3E); the time when the contralateral inhibition reached its maximum (33.10±1.76 ms) matched the time when the spike rate transiently decreased to a local minimum (32.15±0.02 ms) (r=0.96). As this time interval matches the interval between the sound pulses of the calling song, contralateral inhibition enhances the coding of sound pulses and can contribute to reduce spike activity in the pulse intervals of natural chirps (Hedwig, 2016). Reciprocal inhibition of the ON1 neurons was also suggested as a temporal filter for the song pattern (Wiese and Eilts, 1985). Based on our data it may support the processing of sound pulses, but its effect is not sufficient as a mechanism for deciphering the temporal pulse pattern at the thoracic level.

## Acknowledgements

We thank and G. Harrison and S. Ellis for excellent technical support.

## Author Contribution

BH and XZ designed the experimental device, XZ performed the experiments and data analysis, XZ and BH wrote the MS.

## Data availability

Details of the methods described can be obtained from the authors.

